# Photoperiodic induction without light-mediated circadian entrainment in a high arctic resident bird

**DOI:** 10.1101/2020.06.19.160721

**Authors:** Daniel Appenroth, Vebjørn J. Melum, Alexander C. West, Hugues Dardente, David G. Hazlerigg, Gabriela C. Wagner

**Affiliations:** Arctic Chronobiology and Physiology, University of Tromsø, Tromsø, Norway; PRC, INRA, CNRS, IFCE, Université de Tours, 37380 Nouzilly, France

**Keywords:** Photoperiodism, circadian, seasonal reproduction, pars tuberalis, eyes absent, deiodinase, Svalbard ptarmigan

## Abstract

Organisms use changes in photoperiod to anticipate and exploit favourable conditions in a seasonal environment. While species living at temperate latitudes receive day length information as a year-round input, species living in the Arctic may spend as much as two-thirds of the year without experiencing dawn or dusk. This suggests that specialised mechanisms may be required to maintain seasonal synchrony in polar regions.

Svalbard ptarmigan (*Lagopus muta hyperborea*) are resident at 74-81° north latitude. They spend winter in constant darkness (DD) and summer in constant light (LL); extreme photoperiodic conditions under which they do not display overt circadian rhythms.

Here we explored how arctic adaptation in circadian biology affects photoperiodic time measurement in captive Svalbard ptarmigan. For this purpose, DD-adapted birds, showing no circadian behaviour, either remained in prolonged DD, were transferred into a simulated natural photoperiod (SNP) or were transferred directly into LL. Birds transferred from DD to LL exhibited a strong photoperiodic response in terms of activation of the hypothalamic thyrotropin-mediated photoperiodic response pathway. This was assayed through expression of the *Eya3, Tsh*β and deiodinase genes, as well as gonadal development. While transfer to SNP established synchronous diurnal activity patterns, activity in birds transferred from DD to LL showed no evidence of circadian rhythmicity.

These data show that the Svalbard ptarmigan does not require circadian entrainment to develop a photoperiodic response involving conserved molecular elements found in temperate species. Further studies are required to define how exactly arctic adaptation modifies seasonal timer mechanisms.

**Summary statement:** Svalbard ptarmigan show photoperiodic responses when transferred from constant darkness to constant light without circadian entrainment.

## Introduction

Animals in temperate and high latitudes use changes in photoperiod (day length) to anticipate upcoming seasons and adjust physiology and behaviour accordingly. The involvement of circadian clocks in this photoperiodic time measurement was first suggested by Erwin Bünning, who proposed a so-called ‘external coincidence’ mechanism. According to the Bünning hypothesis (Bünning, 1936), organisms express an innate circadian rhythm of photo-inducibility and light exposure coinciding with the photo-inducible phase of this rhythm triggers a photoperiodic response.

In order to test the Bünning hypothesis, experimental approaches based on artificial light exposures, such as night break experiments, have been employed (Bünning, 1936; Elliott et al., 1972; Follett and Sharp, 1969; Follett et al., 1992; Gwinner and Eriksson, 1977; Hamner and Enright, 1967; Pittendrigh, 1972). Night break experiments trigger a long day response by combining a short photoperiod with a nocturnal light pulse that occurs in the photo-inducible phase. Positive results of these experiments across diverse taxonomic groups favour a circadian-based photoperiodic readout mechanisms.

In birds and mammals, photoperiodic effects on reproduction depend on changes in hypothalamic gonadotrophin releasing hormone (GnRH) secretion at the median eminence, and recent evidence points to a coincidence timer mechanism in the adjacent *Pars tuberalis* (PT) as the key upstream control mechanism (Dardente et al., 2010; Hazlerigg and Loudon, 2008; Lincoln et al., 2002; Masumoto et al., 2010; Nakao et al., 2008; Yasuo et al., 2003; Yoshimura et al., 2003).

Within the PT, long photoperiods (LP) stimulate the expression of the thyroid stimulating hormone (TSH) β subunit (*Tsh*β) (Nakao et al., 2008). LP induced expression of TSH leads to increased *Dio2* expression in the mediobasal hypothalamus (MBH), through a cAMP dependent pathway in neighbouring ependymal cells known as tanycytes (Bolborea et al., 2015; Hanon et al., 2008; Nakao et al., 2008; Ono et al., 2008). DIO2 locally converts thyroxine (T_4_) to the bioactive triiodothyronine (T_3_) by outer ring deiodination, thus increasing hypothalamic T_3_ concentration under LP. In long day breeding birds and mammals, this in turn increases the release of GnRH in the median eminence, ultimately leading to gonadal activation (Yamamura et al., 2004; Yamamura et al., 2006; Yoshimura et al., 2003). Conversely, under short photoperiod, low levels of TSH in the PT coincide with increased type III iodothyronine deiodinase (*Dio3*) expression in tanycytes, keeping hypothalamic T_3_ concentration low and promoting gonadal inactivation (Yasuo et al., 2005). The reciprocal regulation of *Dio2*/ *Dio3* expression and the resulting bioactive T_3_ concentration in the MBH is at the core of photoperiodic control of seasonal reproduction and has become a central paradigm in photoperiodic time measurement.

Several lines of evidence suggest that this PT-mediated readout system is circadian-based. First, in both birds and mammals so-called ‘clock genes’ show characteristic rhythmical expression in the PT/ MBH region, consistent with a possible coincidence timer mechanism (Johnston et al., 2005; Lincoln et al., 2002; Tournier et al., 2007; Yasuo et al., 2003; Yasuo et al., 2004). Secondly, in the Japanese quail (*Coturnix japonica*) photoperiodic induction of *Dio2* and downstream physiological responses can be triggered by night break experiments (Yoshimura et al., 2003), implying control through a coincidence timer mechanism.

Further evidence for the circadian basis on the hypothalamic long day response derives from research on eyes absent 3 (EYA3). In mammals, EYA3 has been proposed to act as a transcriptional co-activator at the *Tsh*β gene promoter and analysis of the ovine *Eya3* promoter demonstrated that its expression is controlled by circadian clock genes (Dardente et al., 2010; Masumoto et al., 2010).

Circadian-based models for photoperiodic time measurement place an emphasis on robust circadian cycles of clock gene expression. This raises the question of what happens in species living at arctic latitudes. Light-dark cycles are absent for extended periods of the year and under such circumstances daily rhythmicity in behaviour and endocrinology breaks down completely (Reierth and Stokkan, 1998; Reierth et al., 1999; Stokkan et al., 1994; van Oort et al., 2005; van Oort et al., 2007). Loss of behavioural and endocrine circadian rhythmicity does not necessarily imply loss of circadian-based photoperiodic response circuits, especially in birds where circadian organisation involves multiple circadian oscillators (Cassone, 2014). Moreover, in temperate bird species, lesioning studies resolve behavioural organisation from photoperiodic sensitivity (Binkley et al., 1972; Menaker and Keatts, 1968; Menaker et al., 1970; Rani et al., 2007; Siopes and Wilson, 1974; Wilson, 1991). Nevertheless adaptation to the Arctic might have had a substantial impact on the entire circadian system, which could also affect circadian-based photoperiodic induction. Fibroblast cultures from reindeer show arrhythmic clock gene expression (Lu et al., 2010) and *in-silico* analysis on clock genes revealed mutations that might impact upon circadian rhythm generation (Lin et al., 2019). If arctic animals cannot sustain circadian rhythmicity in the polar day and polar night, this might limit photoperiodic responses through coincidence timing to those phases of the year with a robust light-dark cycle.

To investigate this, we have performed photoperiod manipulations in captive Svalbard ptarmigan (*Lagopus muta hyperborea* Sundevall, 1845), the northernmost resident herbivorous bird species (Fig. 1). Svalbard ptarmigan are highly seasonal in their breeding physiology (Steen and Unander, 1985; Stokkan et al., 1988; Stokkan et al., 1986) and become behaviourally arrhythmic around the solstices (i.e. during the polar night and the polar day) (Reierth and Stokkan, 1998). Similar dampening of melatonin rhythmicity has also been observed (Reierth et al., 1999).

**Figure 1.**
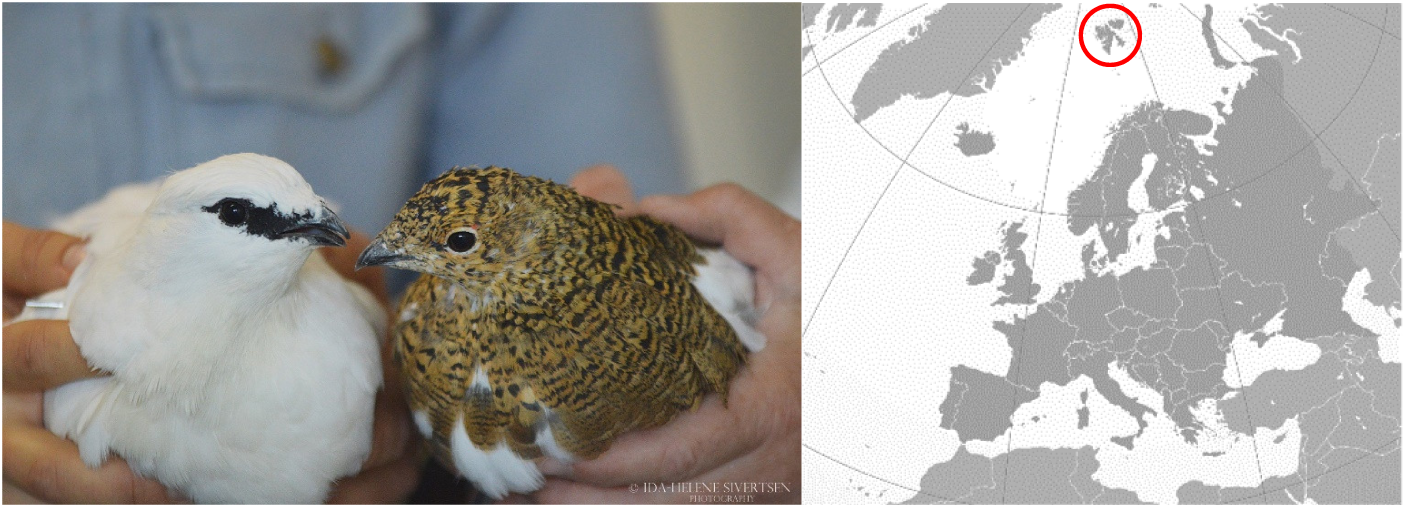
Svalbard ptarmigan (*Lagopus muta hyperborea*) and where to find them. The picture shows a male in white winter plumage and a female in brown summer plumage (Picture taken by Ida-Helene Sivertsen). The Svalbard ptarmigan is a sub-species of the rock ptarmigan (*Lagopus muta*) and inhabits the high arctic archipelago of Svalbard (74° to 81° north latitude).

In order to test if a light-dark cycle is necessary to induce a long day response in Svalbard ptarmigan, we transferred birds, acclimated to constant darkness (DD), either into a gradually increasing photoperiod or directly into constant light (LL). The former group therefore received a rhythmic light-dark cycle while the latter did not. The control group remained in DD. We measured gonadal mass and behavioural activity as well as *Eya3, Tsh*β, *Dio2* and *Dio3* expression in the PT/ MBH region.

## Material and methods

### Experimental animals and housing

All animals were kept in accordance of the EU directive 201/63/EU under a licence provided by the Norwegian Food Safety authority (Mattilsynet, FOTS 7971). Chicks were hatched from eggs laid by captive adult Svalbard ptarmigan at the University of Tromsø (69° 39’N, 18° 57’E). Hatching took place between June 24^th^ 2017 and August 1^st^ 2017. The chicks were raised either indoors with a photoperiod corresponding to the on- and offset of natural civil twilight in Tromsø or outside on the ground. Upon reaching a body mass of 400 to 500 g, 29 birds (Table S1) were transferred into individual cages (1.5 m x 0.5 m) in light and temperature controlled rooms. All birds were transferred at the end of September 2017. Food (standardised protein food; Norgesfor, Ref. No.:OK 2400 070316) and water were provided *ad libitum* throughout the study. Female and male birds were housed together.

Controlled lighting was provided by fluorescent strip lights (Osram L 58 W 830 Lumilux) delivering approximately 1000 lux at floor level. All rooms were further equipped with permanent red illumination (Philips BR125 IR 250 W). During the initial acclimation phase the photoperiod was gradually decreased until reaching DD (red light excepted) on December 22^nd^ 2017. Birds in DD were held under red light to allow for husbandry. The birds remained in DD for five weeks prior to experimental light treatments.

### Experimental light treatment and sampling

After five weeks of DD five individuals were sampled as an initial control group. This marked the start of the experiment (point 0). Thereafter, the three experimental groups were transferred to their respective light treatments (Fig. 2 and Table S1). Six birds remained in DD until the end of the experiment, nine birds were directly transferred into LL and nine birds were exposed to a simulated natural photoperiod (SNP). The SNP treatment reflected an increase in day length following the progression of civil twilight on- and offset of Longyearbyen, Svalbard (78°13′N 15°38′E; Table S2).

**Figure 2.**
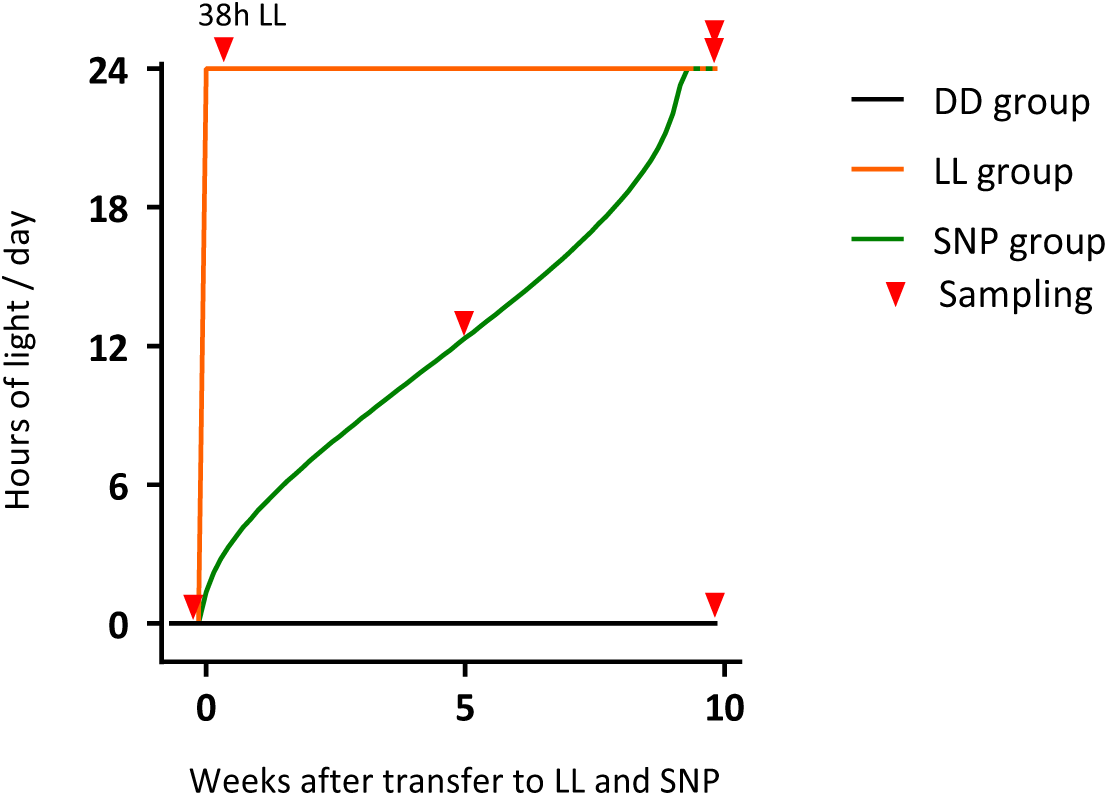
Experimental design. Constant darkness adapted birds were either transferred into constant light (LL group), simulated natural photoperiod (SNP group) or retained under constant darkness (DD group). Red arrows indicate sampling points.

Four individuals were sampled after 38 hours in LL. This sampling time was chosen to coincide with acute photoperiodic gene induction as previously reported in the quail MBH and PT (Nakao et al., 2008). Subsequent samplings aimed to investigate chronic changes in gene expression, and were undertaken at single time points on the following days: After five weeks, four individuals were sampled from the SNP group as they reached LD 12:12. This sampling was performed 3.5 – 4.5 hours after lights on. After ten weeks of light treatment all remaining birds from all groups were sampled. The SNP group had reached LL through a gradual increase in photoperiod four days before the final sampling. All groups were euthanised between 9:00 and 15:00 local time. The DD group was euthanised on the day after the LL and SNP group. Samplings of birds in DD was performed under dim red light only.

Brains were removed after euthanasia and rapidly transferred onto a cooled metal block until stored at −80 °C. Testes and ovaries were removed and measured *post-mortem*.

### Activity

Locomotor activity of all experimental birds was continuously recorded as movement per minute by passive infrared sensors, mounted on the cage doors. Data were collected by an Actimetrics CL200 USB interface coupled to ClockLab data acquisition software (Version 2.61).

### cDNA cloning and *in situ* hybridisation

Probe synthesis and *in situ* hybridisation were performed as described in Lomet *et al*. (2018). RNA was extracted from Svalbard ptarmigan brain tissue using TriReagent (Sigma) and converted into cDNA using Omniscript RT kit (Qiagen). The Icelandic rock ptarmigan genome (Kozma et al., 2016) was used to design PCR primers to amplify cDNA fragments for *Tsh*β, *Eya3, Dio2* and *Dio3*. PCR was performed with Taq DNA polymerase (Qiagen). PCR products of correct sizes were extracted and cloned into pGEMT easy vectors (Promega). The inserts (Table S3) were sequenced (Eurofins Sequencing services, Germany) and verified against the reference genome.

Cloned vectors were stored at −20°C until further use. Prior to hybridisation, vectors were linearised and transcribed using a Promega transcription kit in combination with a ^35^S-UTP isotope (PerkinElmer) to obtain radioactively labelled complementary riboprobes. The riboprobes were purified with illustra MicroSpin G-50 columns (GE healthcare) and incorporation of ^35^S-UTP was measured by a liquid scintillation counter (Triathler multilable tester, Hidex).

Frozen brains were cryosectioned at 20 µm and sections containing PT and MBH were mounted to pre coated adhesion slides (SuperFrost Plus, VWR). Brain sections were fixed in 4 % PFA (0.1 M PB) for 20 minutes at 4 °C and rinsed twice with 0.1 M PB for 5 minutes. Fixed sections were acetylated with 3.75 % v/v of acetic anhydride in 0.1 M triethanolamine buffer (0.05 N NaOH) and rinsed twice with 0.1 M PB for 5 minutes. Sections were subsequently dehydrated with stepwise increasing ethanol solutions (50 %, 70 %, 96 %, 100 % for 3 minutes each) and dried under vacuum for at least 1 hour.

Dried sections were hybridised with 10^6^ cpm of riboprobe per slide in hybridisation buffer (50 % deionised formamide, 10 % dextran sulfate, 1 x Denhardt’s solution, 300 mM NaCl, 10 mM Tris, 10 mM DTT, 1 mM EDTA, 500 µg/ml tRNA). Hybridisation was performed at 56°C overnight. Hybridised sections were washed with 4 x saline sodium citrate (SSC) solutions (3 x 5 minutes) and treated with RNase-A solution (500 mM NaCl, 1 mM Tris, 1 mM EDTA, 20 µg/ml) for 30 minutes at 37 °C. Subsequent stringency washes were performed in SSC (supplemented with 1 mM DTT) of decreasing concentration: 2 x SSC (2 x 5 minutes), 1 x SSC (1 x 10 minutes), 0.5 x SSC (1 x 10 minutes), 0.1 x SSC (30 minutes at 60°C), 0.1 x SSC (rinse). Slides were dehydrated afterwards in stepwise increasing ethanol solutions (50 %, 70 %, 96 %, 100 % for 3 minutes each) and dried under vacuum. Dried sections were exposed to autoradiographic films (Carestream Kodak BioMax MR film) for 9 to 12 days. Exposed films were developed, fixed and digitalised with an Epson transmission scanner. Optical density (OD) was measured with ImageJ (Version 1.51k, Wayne Rasband).

### Analysis

Actograms were produced with the ActogramJ plugin for ImageJ (Schmid et al., 2011) and period length of activity was measured by chi-squared periodograms produced by the same program.

Graphs of gene expressions in the PT/ MBH region and gonadal mass were prepared in GraphPad Prism 8 (Version 8.0.2). The results were plotted as each replicate with lines going through the respective mean of each group at each sampling point. Statistical comparisons were made by 1 way ANOVA and Tukey’s post hoc tests, performed on log transformed values to ensure homogeneity of variances; the threshold for significance was p < 0.05.

Individual values for gene expression with the corresponding gender can be found in Table S1.

## Results

### Activity rhythms

Prior to the experimental treatment, all birds in DD exhibited short episodic bouts of activity with no clear periodicity (Figs 3, S1, S2), and for birds continuing on DD the same pattern was maintained. In birds transferred to LL, episodic activity continued, sometimes with ultradian periodicity. Period lengths were typically in the range 3 – 20 h, and highly variable between individuals. Birds transferred to SNP, based on Svalbard civil twilight progression, showed robust daily rhythms with a period of 24h (p < 0.05).

**Figure 3.**
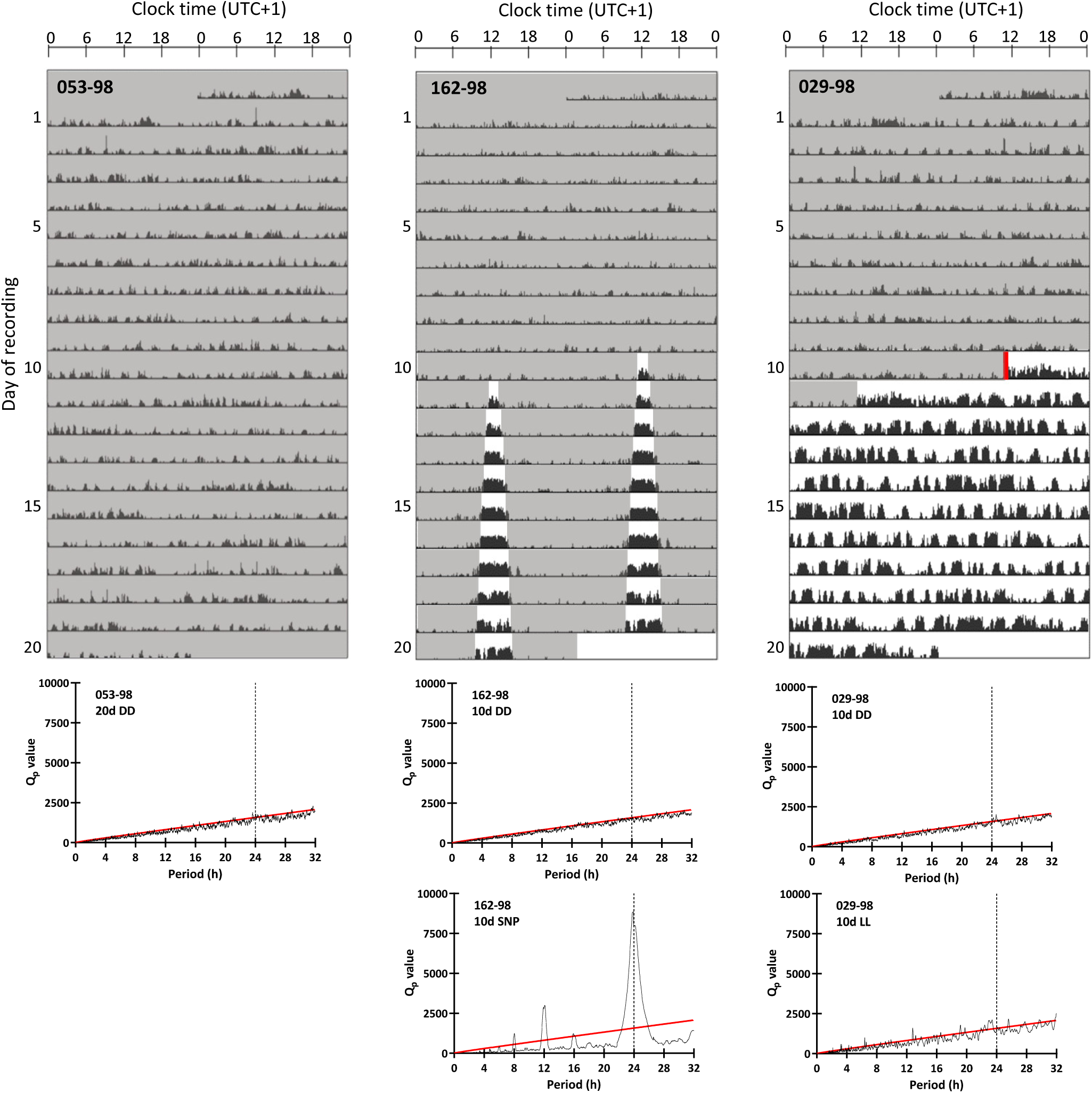
Representative actograms and their respective chi-squared periodograms. Birds adapted to constant darkness were transferred to their respective light treatments on day 10 of the recording (red line) or retained in DD. Actograms are double plotted and grey shadings indicate periods of darkness. Chi squared periodograms were produced for 20 days for the DD group or 10 days before and within experimental photoperiod for the LL and SNP group (upper periodogram: 10d before light treatment (DD), lower periodogram: 10d in light treatment). Q_P_ values above the red line in the periodogram indicate significant periods (p<0.05).

### Gonads

Testes and ovaries were initially regressed in all groups (Fig. 4), and subsequent development depended on photoperiodic treatment (p < 0.0001 by 1 way ANOVA in both cases). Exposure to LL strongly stimulated gonadal maturation for both testes and ovaries, so that after 10 weeks masses increased 22-fold and 93-fold, respectively (p < 0.0001 by Tukey’s post hoc test in both cases). Gonadal maturation in birds maintained in DD and in female birds under SNP, was negligible (DD = 1.4-fold, SNP = 1.1-fold compared to initial values) while male birds transferred to SNP showed a more modest (3.2-fold) but nonetheless statistically significant increase increased testicular mass by the end of the study (p < 0.001 by Tukey’s post hoc test).

**Figure 4.**
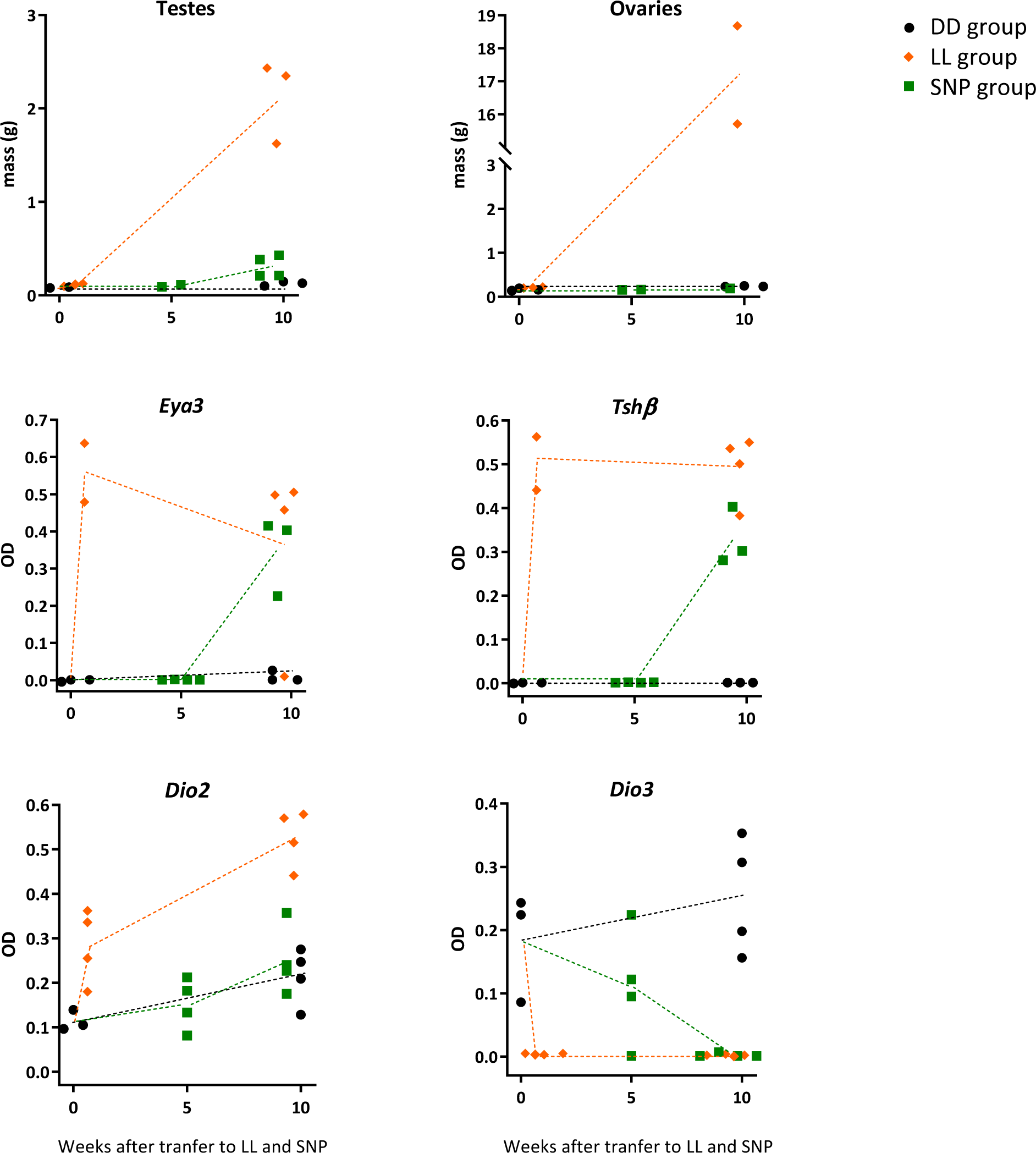
Gonadal development and gene expression in the MBH measured by *in situ* hybridisation. Gonad mass was measured *post-mortem*. Hypothalamic genes were measured before (point 0) and 10 weeks after the transfer into the respective light regime. Additionally gene expression was measured after 38 hour in LL and 5 weeks after the transfer into the simulated natural photoperiod (L:D 12:12).The gene expression is given in optical density (OD) and each replicate is plotted with dotted lines going through the respective mean.

### *Eya3* and *Tsh*β expression

The expression of *Tsh*β and *Eya3* over the course of the study was dependent on photoperiod (p < 0.0001 by 1 way ANOVA in both cases) (Figs 4, 5). Expression of both genes was below the detection threshold at week 0, and rose dramatically 38 hours after the transfer to LL (p < 0.001 in both cases by Tukey’s post hoc test). Thereafter expression of both genes was maintained at high levels until the end of the study (week 10).

**Figure 5.**
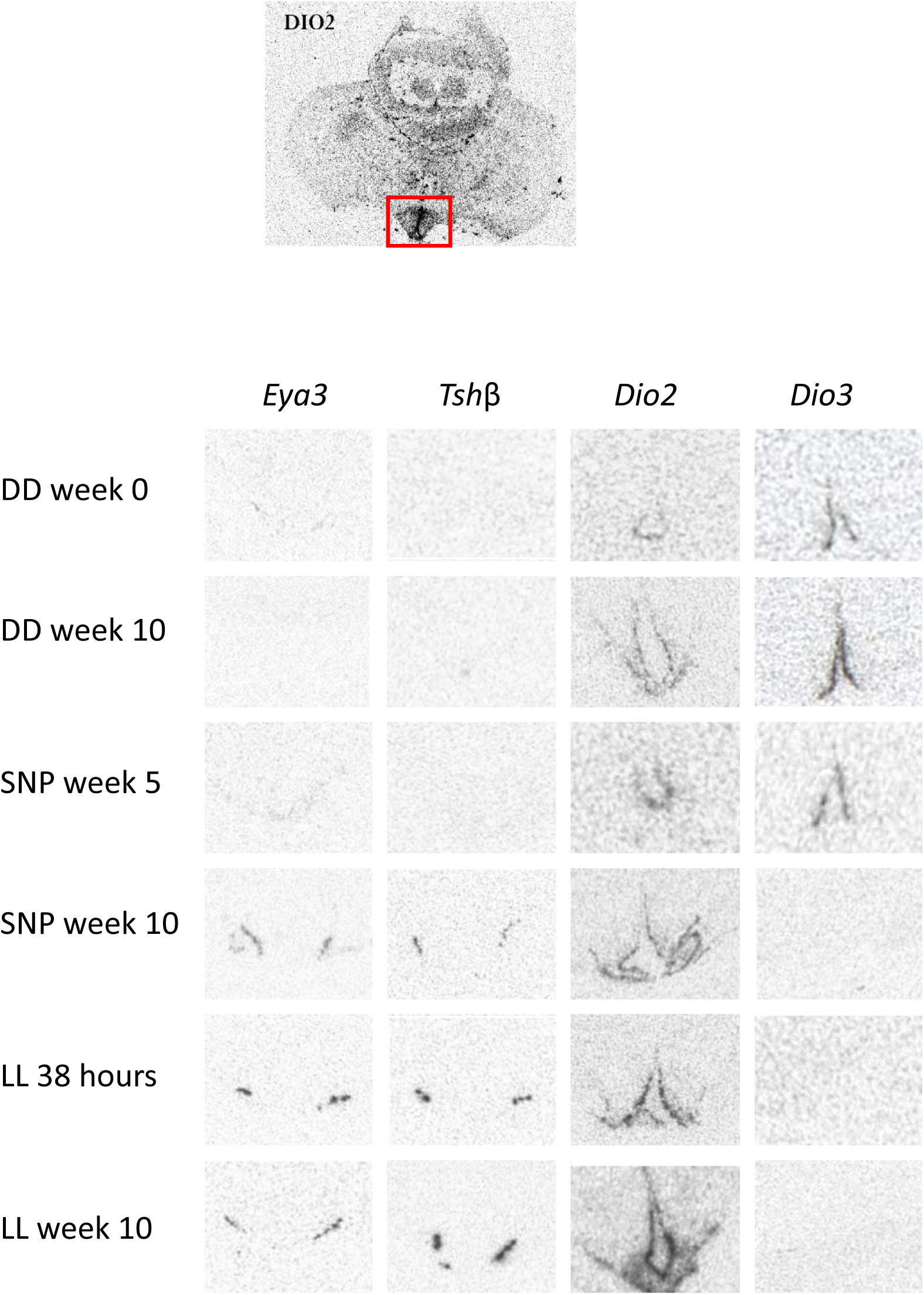
Representative *in situ* hybridisation radiographs for each gene and each sampling point. Top pictures shows whole brain radiograph for *Dio2* highlighting the region of interest (MBH and PT). Radiographs for the respective sampling points show the PT/ MBH region.

In birds exposed to SNP, levels of both genes remained undetectable five weeks after the transfer, when the photoperiod had increased to 12 hours of light. Subsequently, after the photoperiod had progressively increased to LL, expression of both genes increased dramatically to values similar to those in the LL treatment group (p < 0.001 compared to initial values by post hoc Tukey’s test).

In birds maintained on DD, levels of both genes remained basal throughout the experiment.

### *Dio2* and *Dio3* expression

*Dio2* and *Dio3* in the ependymal region of the MBH showed reciprocal changes in expression over the course the study (p < 0.0001 by 1 way ANOVA) (Figs 4, 5). Initial *Dio2* expression was relatively weak, while *Dio3* expression was relatively strong (week 0). Transfer to LL increased *Dio2* expression 2.5-fold within 38 hours (week 0 vs 38 hours LL; p < 0.05 by post hoc Tukey’s test), while over the same period *Dio3* expression was suppressed to background levels (45-fold decrease; p < 0.01 by Tukey’s post hoc test). Under continued LL exposure, elevated *Dio2* levels and suppressed Dio3 levels were maintained to the end of the experiment.

Expression levels of *Dio2* and *Dio3* from birds under SNP gradually increased and decreased respectively over the course the study. In both cases, expression levels after 5 weeks under SNP did not differ from initial values, while levels at week 10 were increased 2.3-fold for *Dio2* and decreased 60-fold for *Dio3* (p < 0.05 and 0.01, respectively by post hoc Tukey’s test).

Under constant darkness, no significant changes in either *Dio2* or *Dio3* expression were observed.

## Discussion

In our experiment we transferred DD acclimated Svalbard ptarmigan either into a simulated natural photoperiod or directly into LL. Both photoperiodic treatments caused increased *Eya3* and *Tsh*β expression and changes in the downstream deiodinases expression but birds transferred from DD to LL displayed no circadian behaviour. This absence of circadian rhythmicity in combination with the lack of an external light-dark cycle might question the circadian basis of the long day response in Svalbard ptarmigan.

According to theory, a circadian-based rhythm of photo-inducibility triggers a photoperiodic response if light exposure occurs during the photoinducible phase (Bünning, 1936). Modern formulations of Bünning’s model focus on events in the PT and the MBH, where night-break protocols induce a long day response in local *Tsh*β expression and downstream effects on hypothalamic deiodinase genes (Dardente et al., 2010; Masumoto et al., 2010; Yoshimura et al., 2003). In sheep, promoter analysis of *Eya3*, a co-activator for *Tsh*β, demonstrates transcriptional control through clock genes, further emphasising the circadian basis for photoperiodic time measurement (Dardente et al., 2010).

Contrastingly, previous studies on arctic animals report the absence of circadian rhythmicity and suggest this as a possible adaptation to polar latitudes, allowing around the clock foraging in constant arctic light conditions (Lin et al., 2019; Lu et al., 2010; Reierth and Stokkan, 1998; Reierth et al., 1999; van Oort et al., 2005; van Oort et al., 2007). Our study confirms the absence of circadian activity rhythms in DD and LL. In a separate experiment we further found no evidence of circadian body temperature rhythms in DD and LL (Appenroth et al, unpublished).

This absence of behavioural and physiological rhythmicity does not exclude the possibility of latent circadian rhythmicity persisting in a coincidence timer mechanism. In non-arctic bird species LL can disrupt circadian activity rhythms but still triggers a photoperiodic response in reproduction (Agarwal et al., 2017; Lumineau and Guyomarc’h, 2003; Simpson and Follett, 1982; Wever, 1980). Moreover, Japanese quail show sustained hypothalamic expression of clock genes in LL, despite behavioural arrhythmicity (Lumineau and Guyomarc’h, 2003; Simpson and Follett, 1982; Yasuo et al., 2003). It therefore remains possible that a sustained rhythm of photo-inducibility may also persist within the PT/ MBH region of arctic Svalbard ptarmigan in constant photic conditions. Consequently the DD-to-LL treatment triggers a long day response as light coincides with the photoinducible phase repeatedly after the transfer (Fig. 6A). Alternatively, the transition from DD to LL might initiate a dampening rhythm of photo-inducibility (Fig. 6B), either by direct induction or by bringing internally desynchronised cellular rhythms into phase (Balsalobre et al., 1998; Nagoshi et al., 2004; Welsh et al., 2004). This scenario would have similar consequences to the persistent rhythmical photo-inducibility described previously and may prove difficult to resolve from one another.

**Figure 6.**
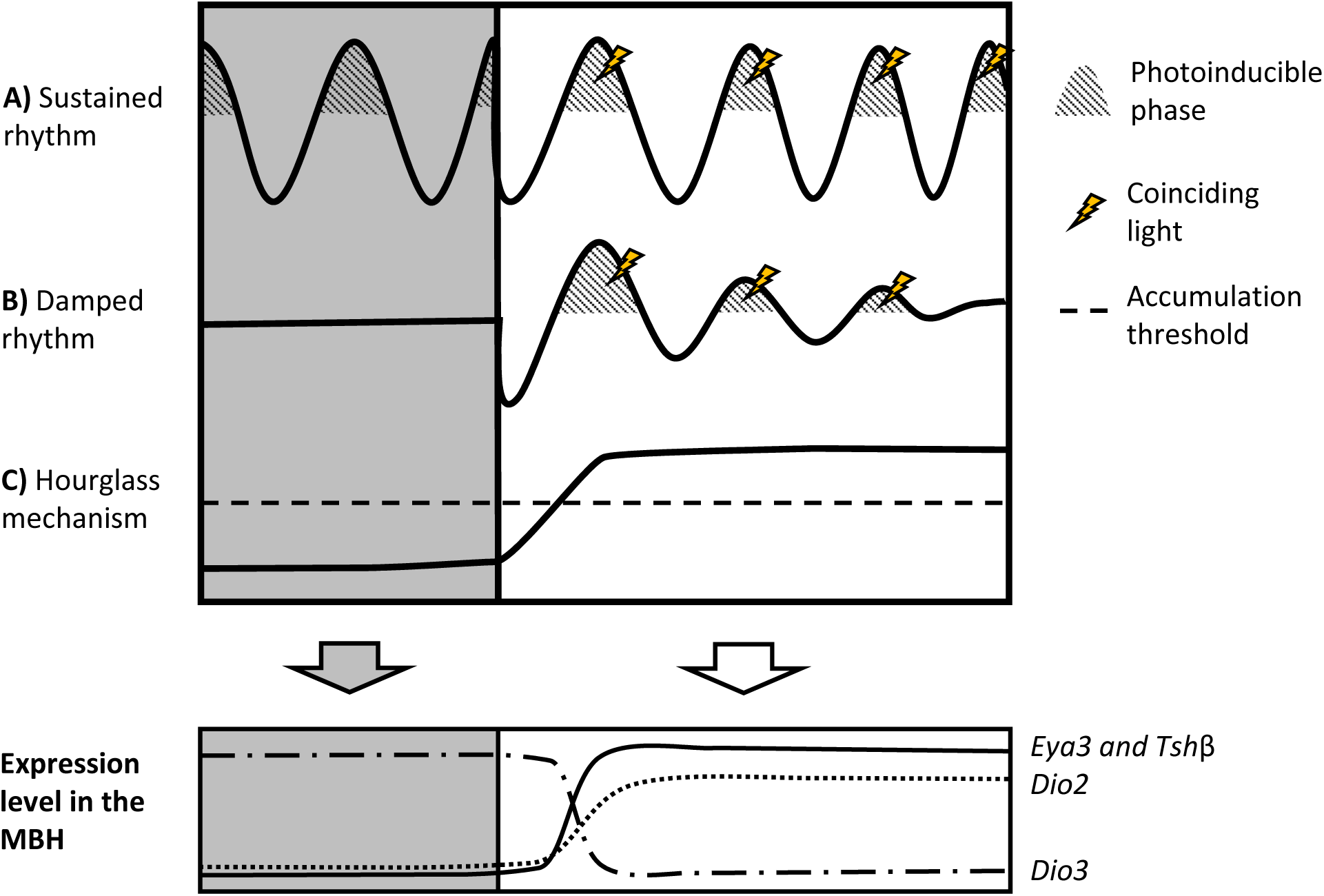
Proposed mechanisms of photoperiodic time measurement in the Arctic. Svalbard ptarmigan show hypothalamic gene expression characteristic for seasonal reproduction when transferred from DD into LL. This process has been proposed to consist of a circadian rhythm of photo-inducibility and coinciding light. Despite absent rhythm in activity a light sensitivity rhythm might be sustained in the PT and MBH throughout constant conditions (A). The rhythm of photo-inducibility might also be initiated by one dawn either by inducing the rhythm or by synchronising individual cells (B). Lastly, the photoperiodic response might be circadian.

Finally, we do not formally exclude that an hour-glass type mechanism operates in these birds. Under this scenario induction relies on the progressive accumulation of a light dependent factor under LL (Fig. 6C). However, we favour a rhythm based model since our molecular characterization of the photoperiodic response shows broad conservation with species known to rely on coincidence timing, like quail (Nakao et al., 2008; Yasuo et al., 2005; Yoshimura et al., 2003) or sheep (Dardente et al., 2010).

Similar to Svalbard ptarmigan transferred from DD to LL, birds subjected to a simulated light-dark cycle showed also increased *Eya3* and *Tsh*β expression and changes in the downstream deiodinases expression at the final sampling point in LL but not earlier in the study when the birds were on L:D12:12. This is consistent with other mammals and birds which require a photoperiod between 12.5 to 14 hours for acute changes of photoperiodic genes in PT and MBH (Hanon et al., 2010; Hanon et al., 2008; Król et al., 2012; Nakao et al., 2008; Ono et al., 2008). By the end of the study, birds in the SNP group showed only limited gonadal development. This is in line with earlier reports that wild Svalbard ptarmigan undergo a delay of several weeks in gonadal development even after exposure to long days (Stokkan et al., 1986).

In summary, our study showed that a high arctic bird relies on the same molecular photoperiodic factors in the PT and MBH to initiate reproduction as other seasonal mammals and birds. Similar responses were measured in birds going through a SNP and birds directly transferred from DD to LL. The latter observation can reasonably be explained by a variant form of coincidence timer mechanism similar to that seen in temperate species. Further experiments using night break or Nanda Hamner protocols (Saunders, 2005) provide a route to test this hypothesis.

## Acknowledgements

The Authors would like to thank the animal technicians from the Arctic Chronobiology and Physiology research group: Hans Lian, Hans-Arne Solvang, and Renate Thorvaldsen. Past, present, and future projects would not be possible without their dedication and experience. We would also like to thank Andreas Nord for all his help with animal handling and the knowledge he shared.

