## Supplementary material for "Photoperiodic induction without light-mediated circadian entrainment in a high arctic resident bird"

**Supplementary information**

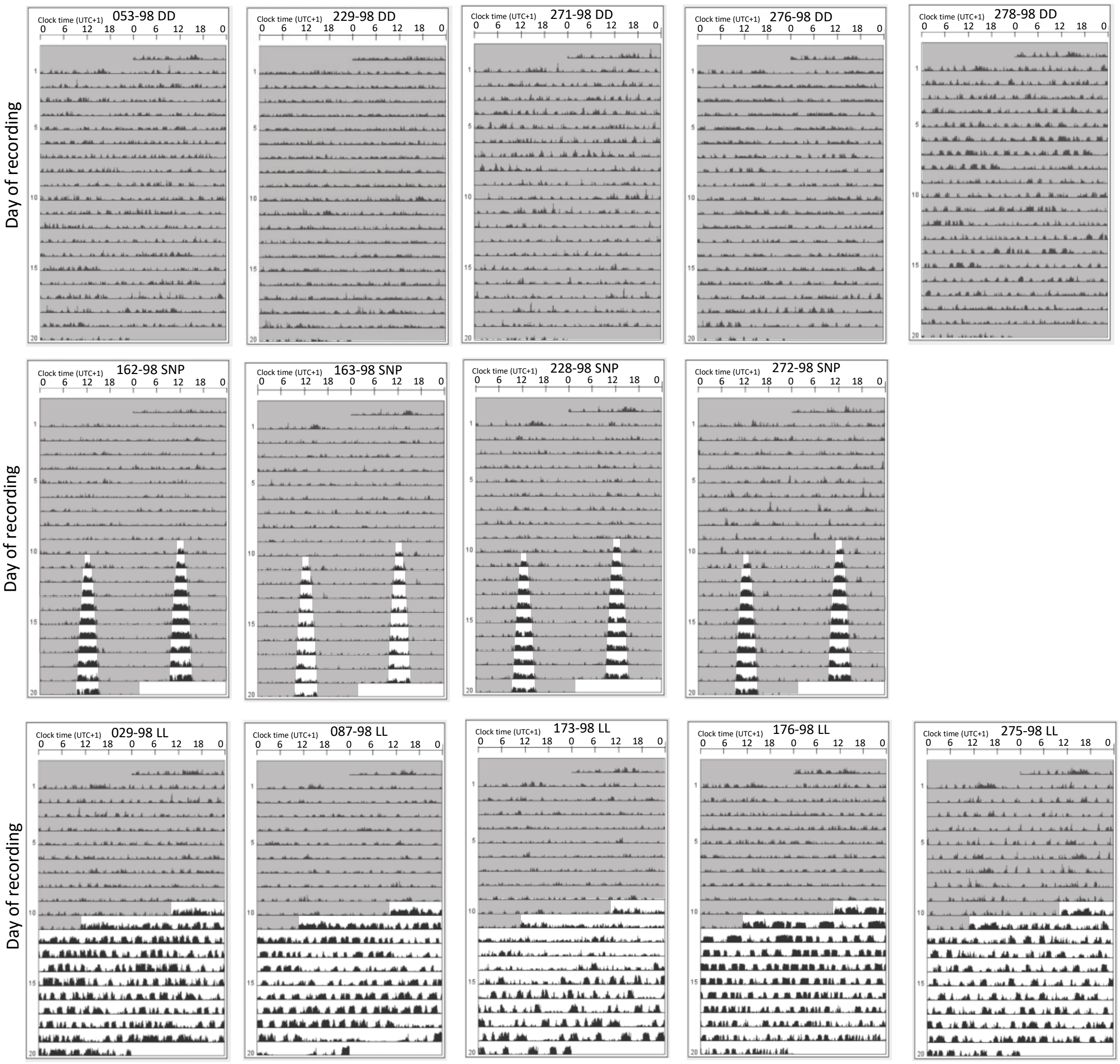


**Figure S1**. **Actograms from experimental birds.** The DD group was plotted for 20 days. SNP and LL birds were plotted 10 days before and 10 days after the transfer into their respective light treatment. Actograms are double plotted and grey shadings indicate periods of darkness.

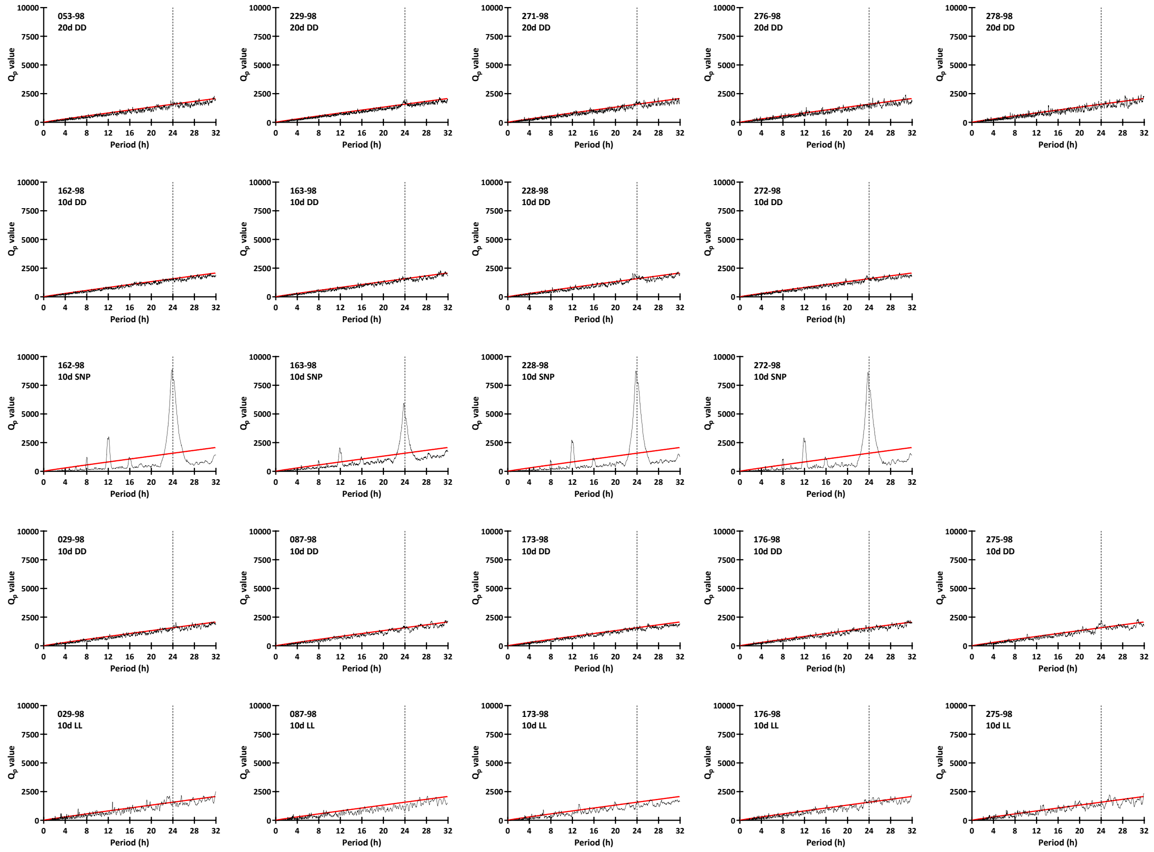


**Figure S2**. **Chi squared periodogram for actograms (Figure S1).** The DD group was analysed for 20 days. SNP and LL birds were analysed 10 days before and 10 days after the transfer into their respective light treatment. $Q_P$ values above the red line in the periodogram indicate significant periods ($p<0.05$).

**Table S1**. **Experimental groups.** Bird identifications and their respective sampling point, gonad mass and optical density (OD) values of hypothalamic gene expression as measured by *in situ* hybridisation. The PT became detached in several brain samples during the sampling procedure. Measurement for *Eya3* and *Tsh*β are therefore missing for several individuals.

| ID | group | gender | Sampling | testes (g) | ovaries (g) | *Eya3* (OD) | *Tsh*β (OD) | *Dio2* (OD) | *Dio3* (OD) |
|---|---|---|---|---|---|---|---|---|---|
| 286-98 | DD | female | Week 0 | | 0.159 | 0.001 | 0.001 | 0.105 | 0.224 |
| 056-98 | DD | male | Week 0 | 0.080 | | 0.001 | 0.001 | 0.096 | 0.243 |
| 037-98 | DD | female | Week 0 | | 0.195 | 0.001 | 0.001 | 0.139 | 0.086 |
| 268-98 | DD | female | Week 0 | | 0.141 | | | | |
| 057-98 | DD | male | Week 0 | 0.087 | | | | | |
| 278-98 | DD | male | Week 10 | 0.129 | | 0.001 | 0.001 | 0.275 | 0.198 |
| 053-98 | DD | male | Week 10 | 0.146 | | 0.001 | 0.001 | 0.247 | 0.156 |
| 229-98 | DD | female | Week 10 | | 0.236 | 0.022 | 0.001 | 0.128 | 0.307 |
| 271-98 | DD | female | Week 10 | | 0.251 | 0.026 | 0.001 | 0.209 | 0.353 |
| 276-98 | DD | male | Week 10 | 0.100 | | | | | |
| 267-98 | DD | female | Week 10 | | 0.234 | | | | |
| 069-98 | SNP | male | Week 5 | 0.113 | | 0.001 | 0.002 | 0.081 | 0.001 |
| 060-98 | SNP | female | Week 5 | | 0.160 | 0.001 | 0.001 | 0.212 | 0.095 |
| 225-98 | SNP | male | Week 5 | 0.090 | | 0.001 | 0.001 | 0.182 | 0.122 |
| 059-98 | SNP | female | Week 5 | | 0.165 | 0.002 | 0.002 | 0.133 | 0.224 |
| 066-98 | SNP | male | Week 10 | 0.382 | | 0.403 | 0.403 | 0.357 | 0.007 |
| 272-98 | SNP | male | Week 10 | 0.207 | | 0.226 | 0.302 | 0.24 | 0.001 |
| 162-98 | SNP | male | Week 10 | 0.428 | | 0.415 | 0.281 | 0.227 | 0.001 |
| 163-98 | SNP | female | Week 10 | | 0.193 | Detached PT | Detached PT | 0.175 | 0.001 |
| 228-98 | SNP | male | Week 10 | 0.211 | | | | | |
| 051-98 | LL | female | 38h in LL | | 0.225 | 0.479 | 0.441 | 0.255 | 0.005 |
| 025-98 | LL | female | 38h in LL | | 0.200 | 0.637 | 0.563 | 0.362 | 0.005 |
| 064-98 | LL | male | 38h in LL | 0.125 | | Detached PT | Detached PT | 0.336 | 0.003 |
| 070-98 | LL | male | 38h in LL | 0.097 | | Detached PT | Detached PT | 0.18 | 0.001 |
| 176-98 | LL | male | Week 10 | 2.434 | | 0.458 | 0.501 | 0.441 | 0.003 |
| 173-98 | LL | female | Week 10 | | 18.681 | 0.498 | 0.536 | 0.57 | 0.002 |
| 275-98 | LL | female | Week 10 | | 15.704 | 0.01 | 0.383 | 0.515 | 0.004 |
| 029-98 | LL | male | Week 10 | 2.349 | | 0.505 | 0.55 | 0.579 | 0.002 |
| 087-98 | LL | male | Week 10 | 1.623 | | | | | |

**Table S2**. **Simulated natural photoperiod (SNP).** The light schedules follows the progression of civil twilight on- and offset in Longyearbyen Svalbard (78°13′N 15°38′E). Clock times are given as coordinated universal time + 1 (UTC+1).

| exp. week | date 2018 | light on | light off | minutes of light/ day | exp. week | date 2018 | light on | light off | minutes of light/ day |
|---|---|---|---|---|---|---|---|---|---|
| | 30.jan | | | 0 | 5 | 07.mar | 06:00 | 18:21 | 741 |
| 0 | 31.jan | 11:31 | 12:53 | 82 | 5 | 08.mar | 05:53 | 18:28 | 755 |
| 0 | 01.feb | 11:06 | 13:17 | 131 | 5 | 09.mar | 05:45 | 18:36 | 771 |
| 0 | 02.feb | 10:48 | 13:36 | 168 | 5 | 10.mar | 05:37 | 18:43 | 786 |
| 0 | 03.feb | 10:33 | 13:51 | 198 | 5 | 11.mar | 05:30 | 18:50 | 800 |
| 0 | 04.feb | 10:20 | 14:04 | 224 | 5 | 12.mar | 05:22 | 18:58 | 816 |
| 0 | 05.feb | 10:08 | 14:17 | 249 | 5 | 13.mar | 05:14 | 19:05 | 831 |
| 0 | 06.feb | 09:57 | 14:28 | 271 | 6 | 14.mar | 05:06 | 19:13 | 847 |
| 1 | 07.feb | 09:46 | 14:39 | 293 | 6 | 15.mar | 04:58 | 19:20 | 862 |
| 1 | 08.feb | 09:36 | 14:49 | 313 | 6 | 16.mar | 04:50 | 19:28 | 878 |
| 1 | 09.feb | 09:27 | 14:59 | 332 | 6 | 17.mar | 04:41 | 19:36 | 895 |
| 1 | 10.feb | 09:17 | 15:08 | 351 | 6 | 18.mar | 04:33 | 19:44 | 911 |
| 1 | 11.feb | 09:08 | 15:18 | 370 | 6 | 19.mar | 04:24 | 19:52 | 928 |
| 1 | 12.feb | 08:59 | 15:26 | 387 | 6 | 20.mar | 04:16 | 20:01 | 945 |
| 1 | 13.feb | 08:51 | 15:35 | 404 | 7 | 21.mar | 04:07 | 20:09 | 962 |
| 2 | 14.feb | 08:42 | 15:44 | 422 | 7 | 22.mar | 03:58 | 20:18 | 980 |
| 2 | 15.feb | 08:34 | 15:52 | 438 | 7 | 23.mar | 03:48 | 20:27 | 999 |
| 2 | 16.feb | 08:26 | 16:00 | 454 | 7 | 24.mar | 03:39 | 20:36 | 1017 |
| 2 | 17.feb | 08:18 | 16:08 | 470 | 7 | 25.mar | 03:29 | 20:46 | 1037 |
| 2 | 18.feb | 08:10 | 16:16 | 486 | 7 | 26.mar | 03:19 | 20:56 | 1057 |
| 2 | 19.feb | 08:02 | 16:24 | 502 | 7 | 27.mar | 03:08 | 21:06 | 1078 |
| 2 | 20.feb | 07:54 | 16:31 | 517 | 8 | 28.mar | 02:58 | 21:17 | 1099 |
| 3 | 21.feb | 07:46 | 16:39 | 533 | 8 | 29.mar | 02:46 | 21:28 | 1122 |
| 3 | 22.feb | 07:39 | 16:46 | 547 | 8 | 30.mar | 02:34 | 21:41 | 1147 |
| 3 | 23.feb | 07:31 | 16:54 | 563 | 8 | 31.mar | 02:21 | 21:54 | 1173 |
| 3 | 24.feb | 07:23 | 17:01 | 578 | 8 | 01.apr | 02:07 | 22:09 | 1202 |
| 3 | 25.feb | 07:16 | 17:09 | 593 | 8 | 02.apr | 01:52 | 22:26 | 1234 |
| 3 | 26.feb | 07:08 | 17:16 | 608 | 8 | 03.apr | 01:34 | 22:47 | 1273 |
| 3 | 27.feb | 07:01 | 17:23 | 622 | 9 | 04.apr | 01:13 | 23:16 | 1323 |
| 4 | 28.feb | 06:53 | 17:30 | 637 | 9 | 05.apr | 00:43 | 00:00 | 1397 |
| 4 | 01.mar | 06:46 | 17:38 | 652 | 9 | 06.apr | | | 1440 |
| 4 | 02.mar | 06:38 | 17:45 | 667 | 9 | 07.apr | | | 1440 |
| 4 | 03.mar | 06:31 | 17:52 | 681 | 9 | 08.apr | | | 1440 |
| 4 | 04.mar | 06:23 | 17:59 | 696 | 9 | 09.apr | | | 1440 |
| 4 | 05.mar | 06:16 | 18:06 | 710 | 9 | 10.apr | | | 1440 |
| 4 | 06.mar | 06:08 | 18:14 | 726 | 10 | 11.apr | | | 1440 |

**Table S3**. **Nucleotide sequences for riboprobe transcription**. Svalbard ptarmigan specific cDNA was cloned into a pGEMT easy vector and the anti-sense riboprobe was obtained by transcription with either SP6 or T7 RNA polymerase (Promega). Sequence is given in the T7 transcription direction.

| | **Size of probe (base pairs)** | **Anti-sense probe synthesis** | **Nucleotide sequence in pGEMT vector (T7 transcription)** |
|---|---|---|---|
| *Eya3* | 772 | SP6 | GGAGGATCACAAACCATGCAGACCTTGTGTCCCTTCACCAAGCCCTTGAGTTAGACTTC<br>CTGTAAGAAGCCAGAGCAAAGGTGTGGCTGTGGCTCCTGCAATCTCTCCATCACTGGG<br>GAGGCAAAGATCTCATAAACAACTGGGGACATTTTCAGCTTGATGAAGAAAACATACC<br>TAATGCAAATTGTACAGTAGACTTCAGGTCATTTCCTCAGGAGCACAGACCTGGCCAAG<br>TCCAGAGGTGTCCTATGAGAAGCACTAGACTCAGCAGAGCTAGAGCTTGACTGGTGAA<br>GAGACTCATGGAATCCAGCTGAATTCTTCTGGCAGGTAGTCTCCAGAGGGAGGGGGTA<br>ACACAGCTGAGAAGGCTCTTACTGATACTTCTGCTTTTCTATTCTGCTTAGTTATAGAAA<br>CCCAAGTAAACATAAAACCTTATTTTTATAGAAAAAAACTGATGGCAGAGCTGAACCTC<br>CCTTGTTTTGCAAAGCCAAAAAGAGATATTGGTTGTTTTTTGTTTGTTTTTTTTTCCATGG<br>GAAATATTAATGAAAATGTCAAAAATACCTCTGCTGCTGTGAAAATGTGTCCTCTCTCCT<br>TCTCTGGGTGTTCAAAAGCAGTTAATTTATTATGATATCCTTACATTATTTCTACAACATG<br>GGATTTTTCCCATTCTGGGATAAGTGGTTCTTAGTAGAGGGAGGTGTTGCTGTTGGTTT<br>CTCCCATTGGCTTCTAGGCAGTGTGTGTAGGATGCACGTGGCTGAGCCCTGTTAATGAG<br>CAG |
| *Tsh*β | 404 | SP6 | TGTCTCTCCTCTTTGGCCTGACCTTTGGTCAAACAGCATCACTTTGTGCTCCTTCAGAGT<br>ACACAATCCACGTGGAGAAACGGGAGTGCGCCTATTGCCTGGCCATCAACACCACCAT<br>CTGCGCCGGATTCTGCATGACTCGGGACAGCAATGGCAAGAAGCTGCTACTCAAAAGT<br>GCTCTGTCCCAAAACGTGTGCACATATAAAGAGATGTTGTATCAAACAGCACTGATTCC<br>GGGCTGTCCTCATCACACCATCCCTTACTATTCCTACCCCGTGGCCATAAGCTGCAAGTG<br>TGGTAAATGTAACACTGACTACAGTGACTGTGTTCACGAGAAGGTTAGGACAAACTAC<br>TGCACTAAGCCACAGAAGCTCTGTAACATGTGAGCTTCCAACAGAACACGG |
| *Dio2* | 693 | T7 | CCAAAGACGAACCTCCTGAAGATTGTAGAAAAAGGGGCCTTTACCTCCCAGGTAGGCA<br>ATTTTTTGTCTCTGCACAATGCATACTCGCTCAAATGAAACCCCATAGGCCACATTGGCA<br>TTGTTGTCCATGCAGTCAGCCACTACTTGGCACTGAGGTGGCAAGGAGAAGTGTTCCA<br>GGAGTTGGTAAGCAGCTGCACATCGATCTTCCTGATTTCTGTGCTTCTTAACTTCAAAG<br>GAAGAGGGGGAGATACCAGGAGCAGCCCAGCCATCTGACGGATGAGCCTCATCGATG<br>TAGACCAACAGAAAGTCAGCCACACCAGAGAACTCTTCCACCAGCTTGCTGAAGGCAG<br>ACAGCTGGCTTGTGAACGGTGGTCAAGTAGCTGAGCCGAAATTAACAACCAGTGGCCG<br>CTCAGAGTTGGCAAAATCCAGAAGATGGCACTTGGCCCCACAGTTACCACCAACACTCT<br>TCCAGCTGCTACTGCTACCATCATTGCGCTTGGCTATGTGGATTACACTGGAATTTGGA<br>GCTTCTCCTCCAAGTTTGACCTGCTTGTAGGCGTCCAGGAGGAAGCTGTTCCAGACGCA<br>GCGCAGCCCCTCCGAGGTCAGCATCCTCCGCCACTCACCGCGCGCAGACTTAGAGCGG<br>CTCAGAAACAGCACCATGTGCTTCAGGAGGATCACAGAGTCATACAGCGC |
| *Dio3* | 556 | SP6 | ATGTTCACGCTGGAGTCGCTGAAGGCTGTGTGGCACGGGCAGAAGCTGGACTTCTTCA<br>AGTCGGCGCACGTGGGCTCGCCGGCCCCCAACCCCGAGGTGATCCAGCTGGACGGGC<br>AGAAGAGGCTCCGCATCCTCGACTTCGCCCGCGGCAAGAGGCCCCTCATCCTCAACTTC<br>GGCAGCTGCACCTGACCCCCGTTCATGGCCCGCCTGAGGTCCTTCCGGCGCCTGGCCGC<br>GCACTTCGTGGACATTGCCGACTTTCTGCTGGTGTACATCGAAGAAGCGCACCCCTCTG<br>ACGGCTGGGTCAGCTCGGACGCTGCCTACAGCATCCCCAAGCACCAGTGCCTCCAGGA<br>CAGGCTGCGGGCAGCGCAGCTGATGCGGGAAGGGGCGCCCGATTGCCCCCTGGCCGT<br>GGACACCATGGACAACGCTTCCAGCGCTGCCTACGGCGCCTACTTCGAGCGGCTCTAC<br>GTCATCCAGGAGGAGAAGGTGATGTACCAGGGAGGCCGAGGACCGGAGGGTTACAA<br>GATCTCGGAGCTGCGGAGCTGGCTAGACCAGTACA |
